# Consistency in macroscopic human brain responses to noisy time-varying visual inputs

**DOI:** 10.1101/645499

**Authors:** Keiichi Kitajo, Takumi Sase, Yoko Mizuno, Hiromichi Suetani

## Abstract

It is an open question as to whether macroscopic human brain responses to repeatedly presented external inputs show consistent patterns across trials. We here provide experimental evidence that human brain responses to noisy time-varying visual inputs, as measured by scalp electroencephalography (EEG), show a signature of consistency. The results indicate that the EEG-recorded responses are robust against fluctuating ongoing activity, and that they respond to visual stimuli in a repeatable manner. This consistency presumably mediates robust information processing in the brain. Moreover, the EEG response waveforms were discriminable between individuals, and were invariant over a number of days within individuals. We reveal that time-varying noisy visual inputs can harness macroscopic brain dynamics and can manifest hidden individual variations.

## Introduction

There is considerable in interest in whether consistent neural responses to repeatedly presented identical external inputs are observable in brain dynamics with ongoing fluctuations. The brain is a complex system composed of a number of nonlinear elements (i.e., neurons) inter-connected with excitatory and inhibitory connections. From the viewpoint of computational neuroscience, it is known that model networks of such complex systems produce irregular and chaotic activity due to a balance of excitatory and inhibitory connections (1, 2). Prior experimental studies have reported fluctuating ongoing activity in the brain, even without specific external stimuli (3, 4). Theoretical studies on chaotic dynamics have revealed that small differences in the initial states of a nonlinear dynamical system can lead to large changes in the responses and instability of the system, even when the system is driven by the same external inputs. In this context, the reliability of spike timing, which is defined as the repeatability of the timing of neuronal spikes when a single neuron is repeatedly driven by a time-varying identical input, is of great interest to neuroscientists. Spike timing should provide “reliable” information, especially if the spikes sequentially produce identical changes in postsynaptic neurons, and this repeatability of timing holds in whole neuronal circuits. Notably, a pioneering study using rat cortical slices showed that single-neuron spikes responding to a repeatedly injected noisy current input showed highly consistent patterns with high temporal precision across trials (5). In addition, a monkey study showed that single neurons in the middle temporal area responding to repeated presentations of the same noisy time-varying motion stimulus exhibited synchronized spike patterns across trials, with high timing precision (6). Moreover, a theoretical study showed that intermittent consistent responses in spike timing should be observable in neural networks with chaotic dynamics (2). Although there is ongoing debate on spike-timing reliability in relation to the rate coding idea (7), these prior studies indicate the existence of reliability in spike timing mediated by the consistent responses of dynamical systems to time-varying inputs.

From a nonlinear dynamical system theory viewpoint, one that is not necessarily concerned with neuronal spikes, this feature is called “consistency”, and is defined as the inter-trial repeatability of response waveforms of a system that is being repeatedly driven by the same time-varying input signal, which can include noise as well as chaotic signals. Consistency has been experimentally demonstrated in laser systems (8), and numerically in the chaotic Lorenz model (9). Reservoir computing, a neural network framework for supervised learning that uses the rich dynamics of recurrent neural networks has attracted much attention (10,11). In the context of reservoir computing, consistency is equivalent to the “echo-state property”, in which a high-dimensional recurrent network system with intrinsic dynamics exhibits consistent outputs against identical driving inputs, regardless of the system’s initial states (10, 12). The echo-state property is important for learning reproducible desired outputs for repeated inputs irrespective of the initial state of the system. Reservoir computing has also attracted attention in computational neuroscience. For instance, the reservoir computing paradigm was used to model the dynamics of prefrontal cortex and cognitive functions (12). Although consistency does not explicitly consider the desired outputs of the nonlinear system, consistency is assumed as a basis for the echo-state property in reservoir computing (13). Consistency is also counterintuitive because it occurs when initial conditions differ, even in chaotic dynamical systems whose responses are sensitive to initial conditions. Consistency is therefore important for achieving robust and reproducible information processing, overcoming the instability that is potentially caused by fluctuations and noise in the brain.

Although local field potential (LFP) recordings and electroencephalography (EEG) are spatially crude methods compared with spike recordings of single neurons, they reflect dynamic changes in the excitability of neuronal ensembles associated with synchronous spike inputs in neuronal circuits. Therefore, consistency in EEG waveforms could be associated with information processing mediated by a number of spike communications. To this end, we investigated whether scalp-EEG-assessed macroscopic human brain responses to an identical noisy visual input showed trial-to-trial consistency on an individual basis.

## Results

### Consistency of EEG responses within individuals

Participants (*n* = 130) were presented with a noisy flickering checkerboard stimulus (Fig. 1) and were required to look passively at the stimulus. The gray-level contrast between adjacent squares was temporally modulated by application of Gaussian white noise with one of five different standard deviations (mean = 0, SD = 16, 32, 48, 64, 80, in 8-bit gray level) and one of two noise realizations for each noise level. Participants were repeatedly presented with each of the noise stimuli for 14 times in a randomized order, and using a 63-channel EEG amplifier we recorded high-density scalp EEG signals while participants either viewed the stimuli or rested for 3 min.

**Figure 1.**
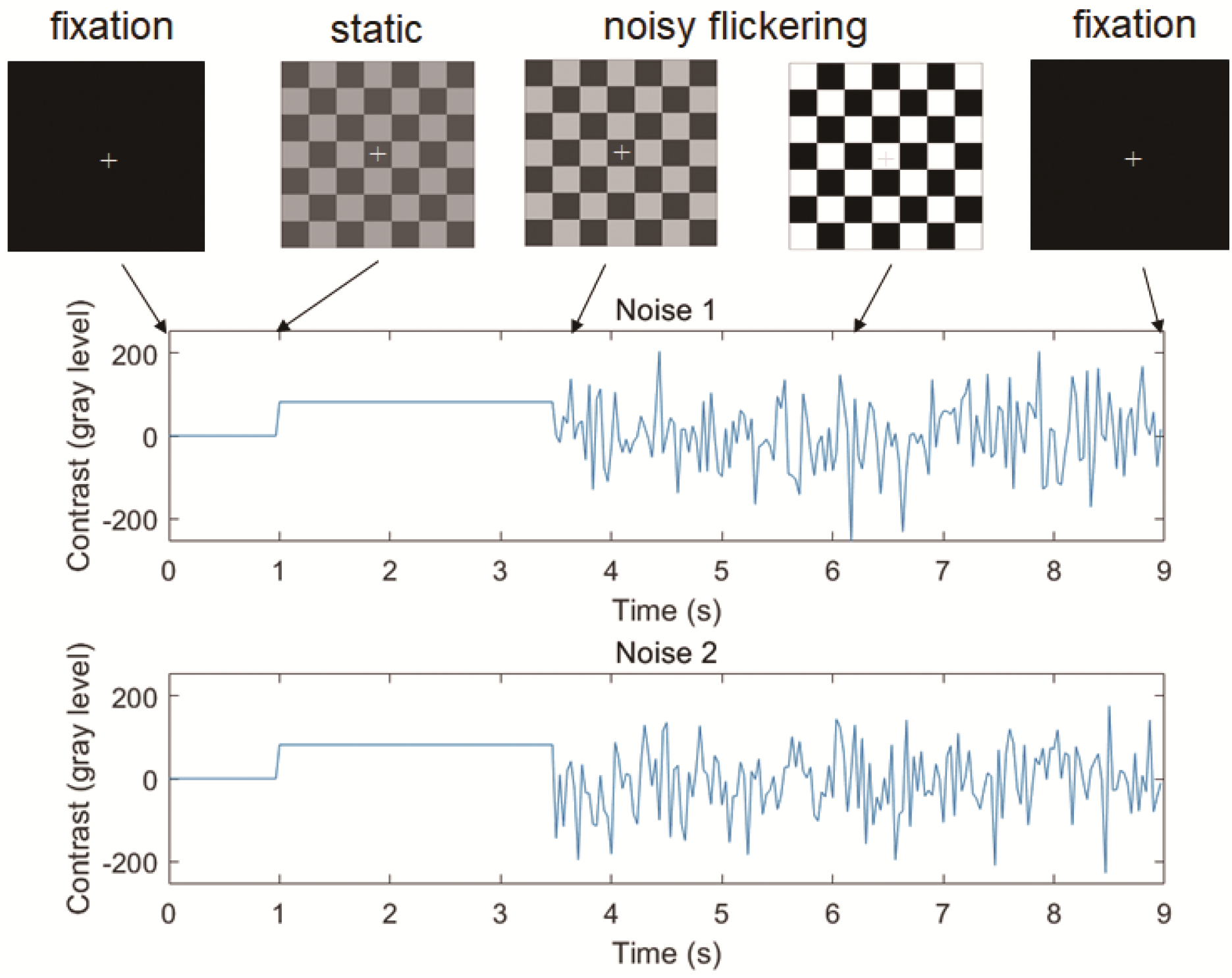
Visual stimuli and experimental paradigm. The stimuli consisted of a fixation cross, a static checkerboard (2.5 s), and a noisy flickering checkerboard (5.5 s).

To assess the degree of consistency of brain responses, we applied a canonical correlation analysis (CCA)-based method between the pairwise EEG epochs within and across individuals. CCA is a conventional statistical method for extracting the linear combinations of data variables that give maximal correlations between pairwise datasets (14), and is useful for detecting synchronization between time series data from two dynamical systems such as two coupled chaotic systems (15). We used CCA to extract correlated variates between pairs of EEG trials, with these correlated variates being considered to reflect consistency in the nature of the brain activity. Fig. 2 indicates the analytical pipeline for the CCA-based method. Specifically, we extracted canonical variates composed of linear combinations of EEG signals and computed the pairwise L1 norm between EEG trials. As there were 14 trials for each of the visual stimuli, we obtained a 28 × 28 L1 distance matrix for each of the 130 participants for each noise intensity level. Next, we applied classical multidimensional scaling (MDS) to analyze the Mahalanobis distance between the centroids of two classes (noise presentation 1 vs 2) and the classification accuracy of a linear support vector machine (SVM) (16) with leave-one-out cross validation (LOOCV) in two-dimensional MDS space.

**Figure 2.**
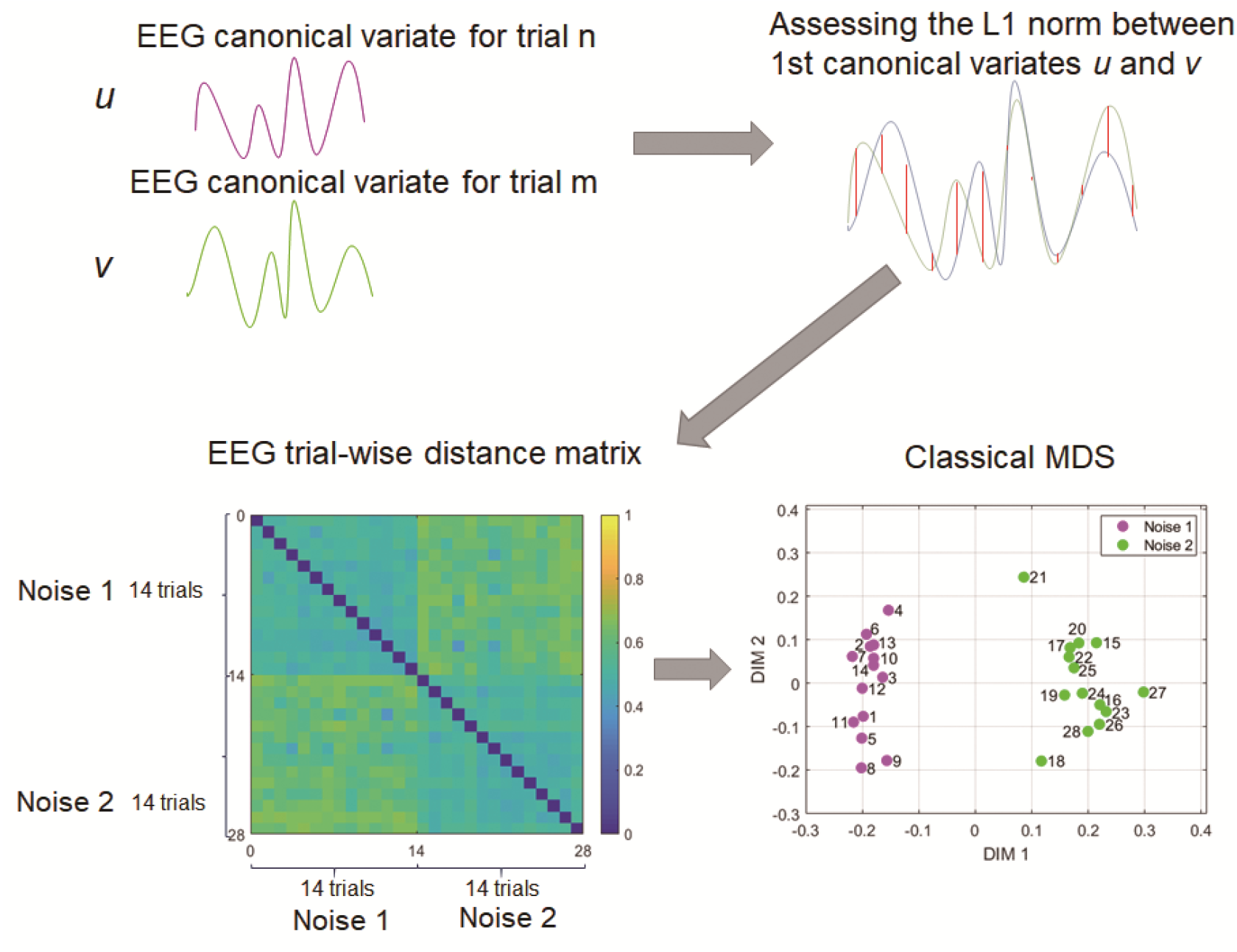
Schematic presentation of the analytical pipeline for the intra-individual analysis. Classical multidimensional scaling (MDS) was conducted using a distance matrix composed of 28 × 28 L1 norm values between EEG canonical variates.

Fig. 3 demonstrates representative 1st canonical variates extracted from two EEG epochs from a single participant for identical (Fig. 3A) and different flickering presentations (Fig. 3B). Although the CCA-based method will try to extract correlated variates from any dataset, the 1st canonical variates derived from a pair of EEG epochs corresponding to the same presentation of visual noise show a much higher canonical correlation than a pair of epochs corresponding to different presentations of visual noise. The 1st canonical variates extracted from the paired EEG epochs for the same visual noise presentations showed peaks in the 4–8 Hz theta band (Fig. S1). Fig. 3B indicates group data for canonical correlation coefficients between the paired EEG epochs for the same and different visual noise presentations averaged across all 130 participants. We observed that the noise level had significant effects (Friedman test, *F_r_* (5, 645) = 375.59, *p* < 0.001) on the canonical correlations. Post-hoc multiple comparison tests showed that the condition with noise of SD 80 showed higher canonical correlations than the other five lower noise levels for identical visual noise presentations (Wilcoxon signed-rank test, two-sided, Bonferroni-corrected *p* < 0.001). Canonical correlations for the EEG trials for two distinct noise presentations also differed across noise levels (Friedman test, *F_r_*(5, 645) = 33.31, *p* < 0.001; Fig. 4B). In addition, the canonical correlations from EEG trials with the same noise inputs were statistically higher than those for different noise inputs for the five noise levels (noise SD: 16, 32, 48, 64, 80; Wilcoxon signed-rank test, two-sided, Bonferroni-corrected *p* < 0.001). Next, to investigate topographical patterns in the extent of contributions of EEG signals to the canonical variates, we assessed the canonical loading, which is the correlation between projected canonical variates and EEG signals. Fig. 3D shows the topography of absolute values of canonical loading averaged across all participants for noise with an SD of 80. We found prominent signal contributions from occipital electrodes placed over the lower visual cortex. These results suggest that this consistency phenomenon mainly occurs in the lower visual areas, and that the CCA-based method did not extract merely spurious correlations between the paired EEG epochs for the same noise presentations by overfitting the EEG signals from all electrodes.

**Figure 3.**
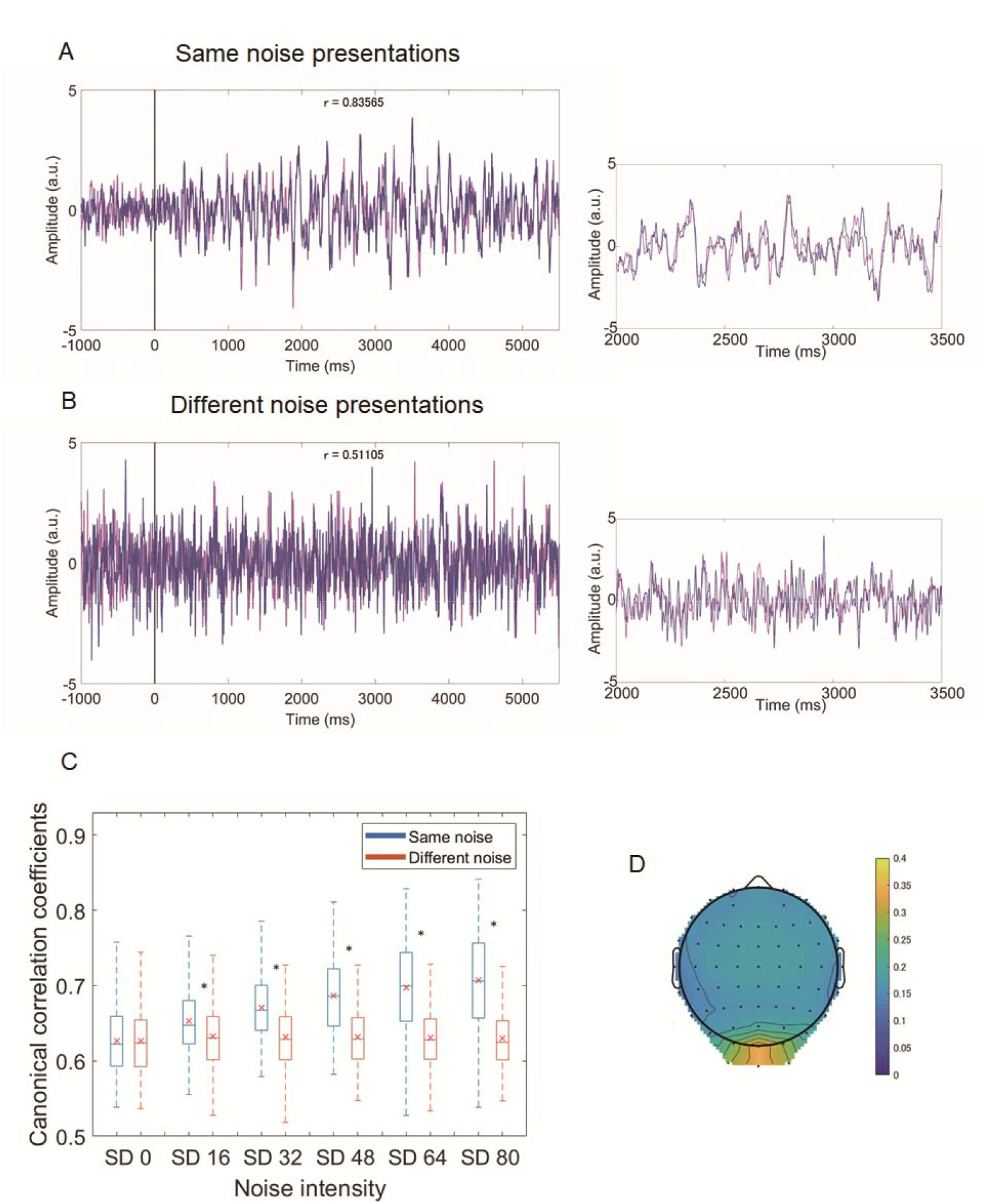
EEG canonical variates, canonical correlation coefficients, and canonical loadings. (A) A pair of EEG canonical variates (magenta and purple lines) extracted from a pair of trials where the participant viewed the same presentation of visual noise. (B) Another pair of EEG canonical variates extracted from trials with different presentations of noise. Visual flicker starts at 0 ms. Right panels are magnified views of the corresponding EEG canonical variates from 2000 to 3500 ms. (C) Group data for canonical correlation coefficients for EEG trials for the same and different noise presentations as a function of noise intensity. Boxplots show the median, 25% and 75% quartiles (boxes), 1.5 times the interquartile range (whiskers), and mean (x). Asterisks indicate significant differences between same and different noise presentations. (D) The topography of absolute values of canonical loadings averaged across all participants (n = 130) for the noise SD 80 condition. Higher loadings were observed for occipital electrodes over the lower visual area.

**Figure 4.**
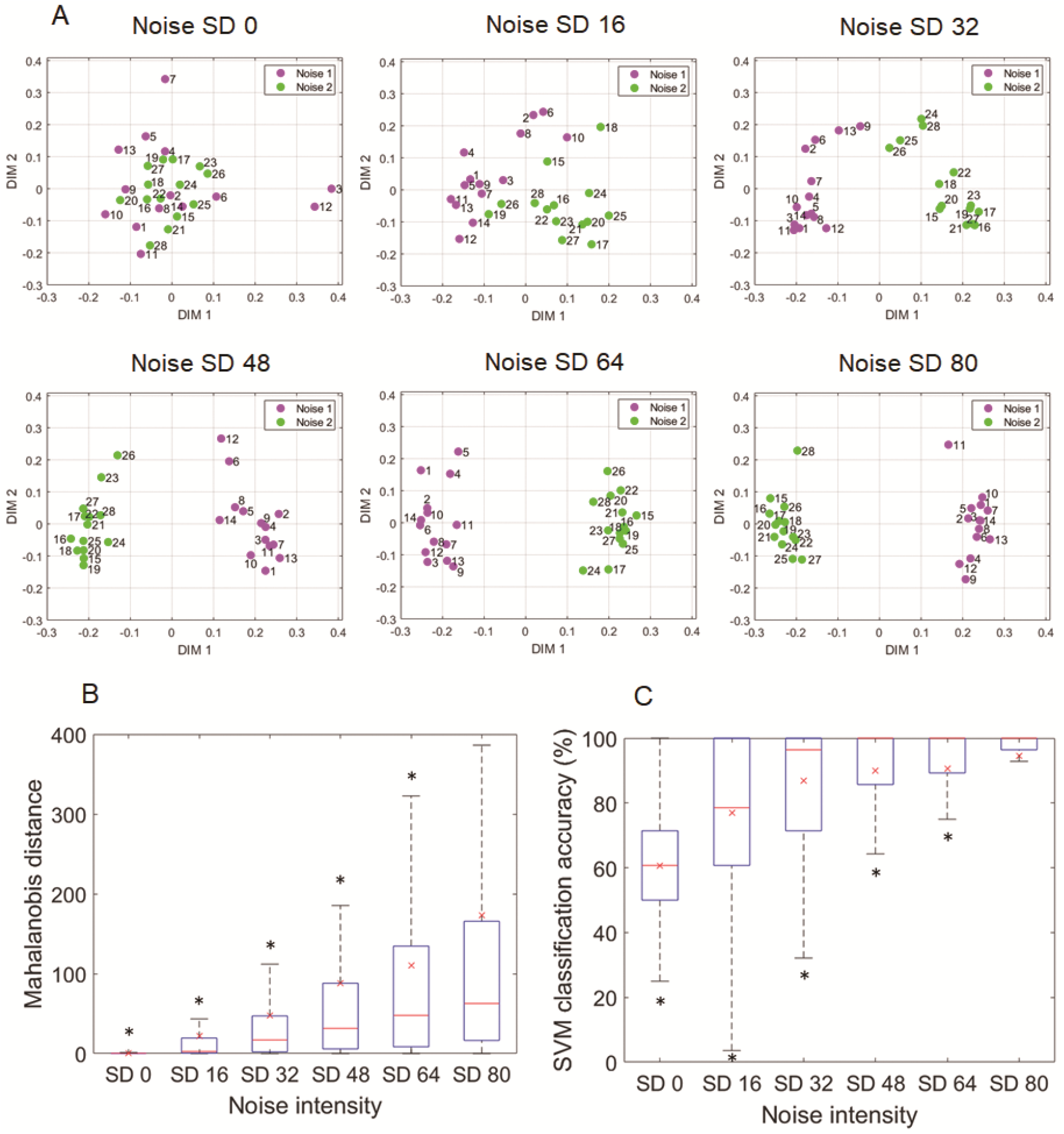
Intra-individual separation of EEG responses corresponding to two noise presentations. **(A)** Effects of noise intensity on separation of EEG trials for two distinct noise presentations. Data from a representative participant. Markers in different colors correspond to different presentations of noise. The numbers indicate the trial number for noise presentations 1 (1–14) and 2 (15–28). **(B)** Group data for the Mahalanobis distance between EEG responses for two noise presentations as a function of noise intensity (n = 130). **(C)** Group data for SVM classification accuracy in MDS space as a function of noise intensity (n = 130). Box plots in **(B)** and **(C)** show the median, 25% and 75% quartiles (boxes), 1.5 times the interquartile range (whiskers), and mean (x). Asterisks indicate significant differences between SD 80 and other conditions.

Next, we analyzed trial-to-trial distance matrices estimated as the L1 norm of two canonical variates and analyzed how EEG trials for distinct noise presentations were located in classical MDS space. Fig. 4A shows an MDS visualization of EEG responses from a representative participant at different noise levels. We found a clear separation between EEG trials from two different visual noise inputs in high noise intensity conditions. To assess the similarity of EEG responses to identical visual inputs in comparison with those to different visual inputs, we estimated the Mahalanobis distance between the centroids of EEG trials for two distinct noise presentations in all participants, with this being the ratio of within-label (noise) variance to across-label (noise) variance in the two-dimensional MDS space. Fig. 4B shows group results for Mahalanobis distances for EEG trials for two distinct visual noise conditions at all noise levels. The Mahalanobis distance monotonically increased as a function of noise intensity, and we observed significant effects of noise intensity on the Mahalanobis distance (Friedman test, *F_r_* (5, 645) = 422.5 *p* < 0.001). Post-hoc multiple comparison tests showed that noise with an SD of 80 showed higher Mahalanobis distances between two noise presentations than did the other noise levels (Fig. 4B; Wilcoxon signed-rank test, two-sided, Bonferroni-corrected, *p* < 0.001). We also used a linear SVM with LOOCV in the classical MDS space to analyze the classification performance of EEG trials corresponding to two visual stimuli. The classification accuracy increased as a function of noise, up to a median correct rate of 100% for noise of SD 48, 64, and 80, and we found that the noise intensity had significant effects on the LOOCV accuracy (Friedman test, *F_r_* (5, 645) = 305.1812, *p* = 0.001; Fig. 4B). Post-hoc multiple comparison tests showed that noise with an SD of 80 showed higher LOOCV accuracy than all other noise levels (Wilcoxon signed-rank test, two-sided, Bonferroni-corrected *p* < 0.001; Fig. 4C). These results indicate that noise with higher intensity showed clear separation of EEG trials corresponding to two different noise presentations, and high classification performance on a single-trial basis.

Taken together, the intra-individual analyses indicate that EEG responses to identical noisy visual inputs show trial-to-trial consistency with the response waveforms differing depending on the input signals, with higher noise levels inducing more prominent consistency. To our knowledge, this is the first evidence that human EEG-level neural signals show a consistency signature in response to noisy visual inputs, other than that from our earlier preliminary results on a small dataset (three participants) (17).

### Inter-individual differences in noise-induced EEG responses

Another intriguing question is whether individual brains, which should differ in various aspects such as network architecture and dynamical characteristics of neurons, show distinct EEG responses to identical noise inputs. To this end, we analyzed individual differences in EEG responses by conducting inter-individual CCAs for identical visual inputs (Fig. 5A). Fig. 5B shows an MDS visualization of EEG responses for a pair of participants at different noise levels. Separation of the individuals was most prominent for the highest noise level. Using the Mahalanobis distance, we estimated the separation of pairwise individuals in two-dimensional MDS space. We found that the noise level had significant effects on the Mahalanobis distance between the centroids of EEG trials for different individuals (Friedman test, *F_r_* (5, 41920) = 33222, *p* < 0.001). Post-hoc multiple comparison tests revealed that the Mahalanobis distance between participants was largest for the highest noise intensity (Fig. 5C; Wilcoxon signed-rank test, two-sided, Bonferroni-corrected *p* < 0.001). We also tested if an SVM could classify EEG trials from different individuals using the LOOCV method, and observed significant effects of noise intensity on the classification accuracy (Friedman test, *F_r_* (5, 41920) = 26690, *p* < 0.001). The SVM classifier showed a median LOOCV accuracy of 100% at the higher noise levels (SD 32, 48, 64, 80), with the highest noise showing higher LOOCV accuracy than all other lower noise conditions (Wilcoxon signed-rank test, two-sided, Bonferroni-corrected, *p* < 0.001, Fig. 5D). These results indicate that EEG responses to identical noise differed across participants. Moreover, in 20 additional participants, the degree of inter-individual separation was higher in the noise SD 80 condition than it was with periodic stimuli (3.75 Hz, 5.0 Hz, 7.5 Hz, 15.0 Hz; Fig. S2).

**Figure 5.**
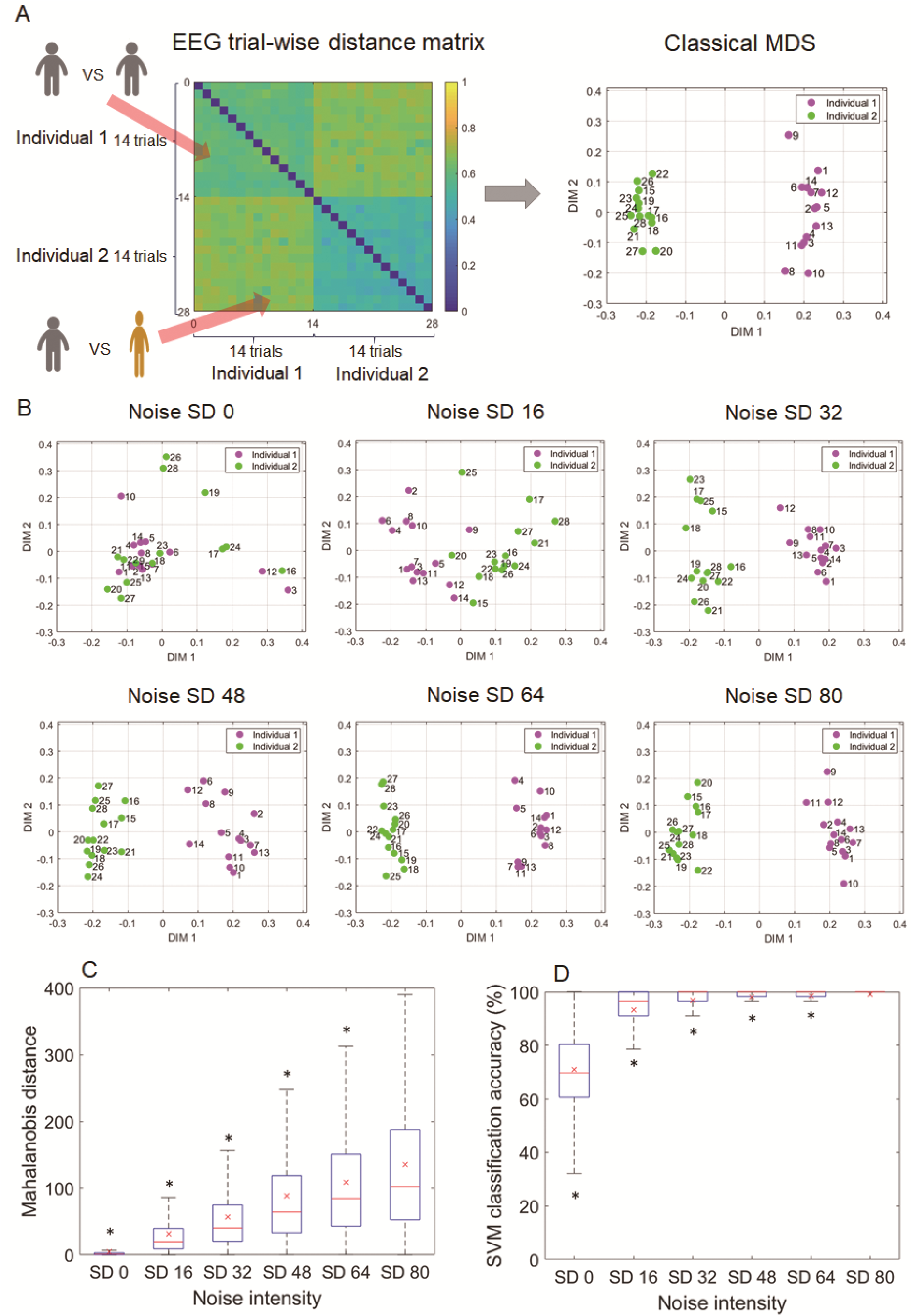
Inter-individual separation of EEG responses. **(A)** Schematic presentation of the distance matrix and MDS visualization. **(B)** Effects of noise intensity on separation of EEG trials for two distinct individuals for presentations of the same noise. Data are from a pair of representative participants for distinct noise levels. The numbers indicate the trial number for individual 1 (1–14) and 2 (15–28). **(C)** Group data for the Mahalanobis distance obtained from pairs of the 130 participants for different visual noise intensities (n = 8385 [= _130_C_2_] participant pairs). **(D)** Group data for SVM classification accuracy in MDS space as a function of noise intensity (n = 8385 [= _130_C_2_] participant pairs). Box plots in **(C)** and **(D)** show median, 25% and 75% quartiles (boxes), range (whiskers), and mean (x). Asterisks indicate significant differences between SD 80 and other conditions.

### Invariance of noise-induced EEG responses in follow-up sessions

We conducted a follow-up testing session in 32 participants at a mean interval of 101 days, and using the individual-wise Mahalanobis distance and the distance rankings of the data from their first recordings, tested whether the second follow-up recordings showed a shorter Mahalanobis distance to the EEG responses of their first recordings than they did to those of the other 129 individuals. Critically, the top 1-ranked (shortest distance) participants were 28 out of 32 participants (87.5%), and top 5-ranked participants were 32 participants among 32 participants (100%) in the individual-wise identification. Fig. 6 shows a t-distributed stochastic neighbor embedding (t-SNE) (18) visualization of the EEG responses of participants. The normalized Mahalanobis distance to other individuals was used as the individual feature for the t-SNE. It can be observed that most of the second recordings of the 32 follow-up participants are located close to their first recordings. These results indicate that within-individual noise-induced EEG responses are invariant across days and show high accuracy in the identification of individuals.

**Figure 6.**
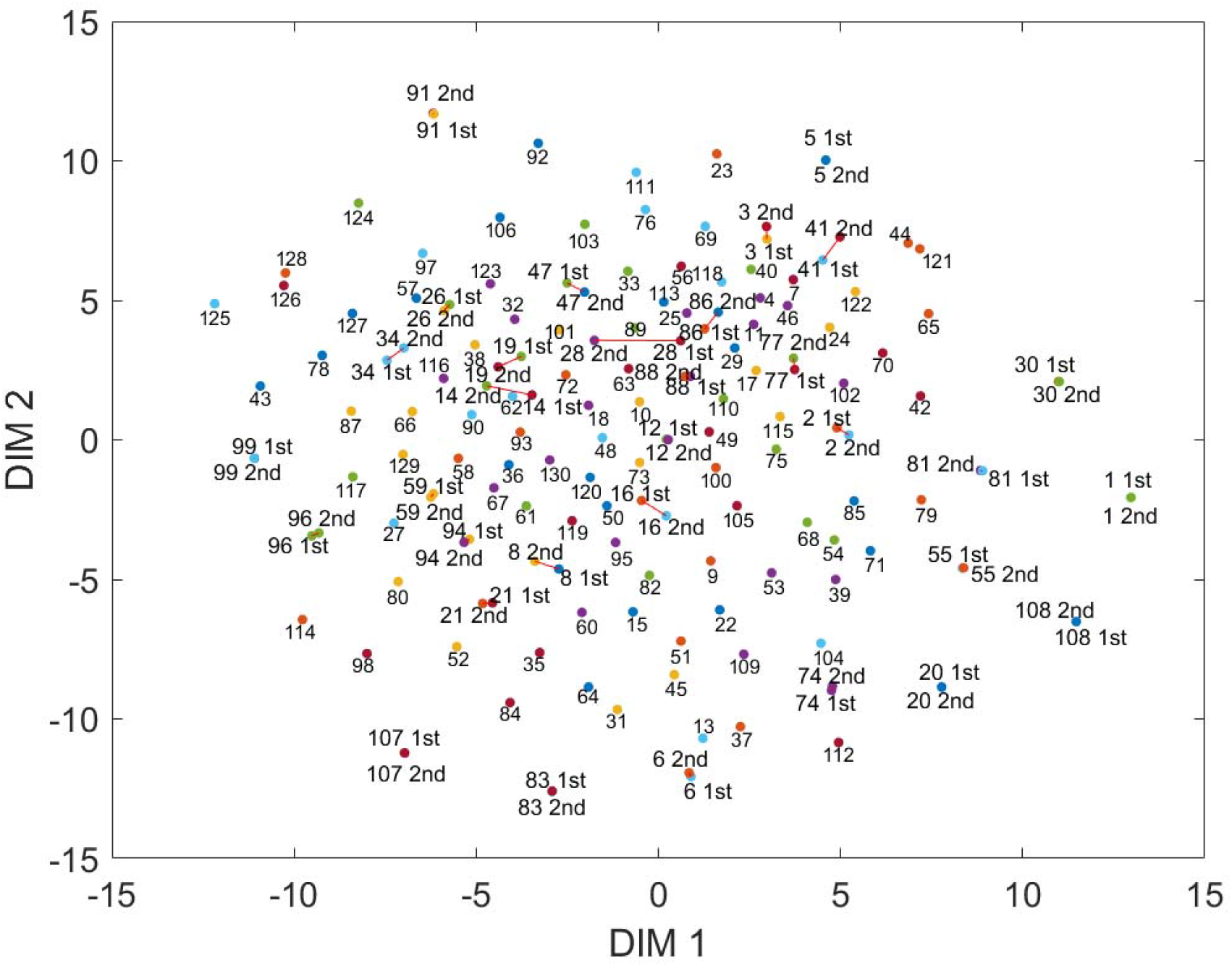
t-distributed stochastic neighbor embedding (t-SNE) visualization of the 130 initial individuals and 32 follow-up individuals. Individuals who underwent follow-up sessions are denoted using 1st and 2nd, in addition to the individual number (1–130), and connections between 1st and 2nd sessions are made with red lines, although most of the follow-up points overlap in t-SNE space (perplexity: 30, exaggeration: 5).

## Discussion

Taken together, large-scale human neural activity exhibits a consistency nature in response to repeatedly presented noisy visual inputs. Scalp EEG is the filtered summation of postsynaptic membrane potentials from a large number of cortical neurons, and a number of studies have shown that a variety of EEG dynamics, such as oscillations and synchrony, are important in mediating perceptual and cognitive processes associated with the spike communications underlying these processes. For example, Fries (19) proposed that coherence in membrane potentials between neuronal groups could constrain spike communication across neuronal groups. We therefore speculate that consistency in the EEG responses to time-varying inputs could be a fundamental basis for robust and reproducible information processing in macroscopic neural activity, which can constrain spike communication in an input-dependent manner.

Differences in initial brain states caused by moment-to-moment fluctuations in ongoing activity could be a potential risk for stable neural responses if the fluctuations were uncontrolled and unrelated to the signals coding information. Animal studies have shown that ongoing cortical activity exhibits a rich and high spatiotemporal variability, even under anesthesia (3,4), and that it modulates evoked responses (3). Non-additive interaction between task-induced and ongoing BOLD activity has also been reported in humans (20, 21), and moment-to-moment variability in BOLD activity is associated with functional performance (22). Additionally, perceptual performance is affected by instantaneous ongoing activity, such as the phase of ongoing EEG oscillations (23, 24). These previous studies suggest that ongoing activity could modulate responses to external stimuli and perceptual performance. Consequently, we speculate that the consistency mechanism observed in the current study reconciles the fluctuating nature of ongoing brain activity with robust and reproducible information processing.

We also demonstrated that noise-induced EEG response waveforms were discriminable between individuals. We speculate that inter-individual differences in network architecture and dynamical characteristics of neurons are associated with distinct responses across individuals. It should also be noted that the individual features could still be detected an average of 3 months later, suggesting that they are robust and associated with individuality. It is also intriguing that the neural code for time-varying visual signals triggered by such a simple checkerboard structure showed individual differences.

It is an open question as to whether these noise-induced nonlinear phenomena play computational roles mediating information processing in the brain. From the viewpoint of nonlinear science, a variety of nontrivial noise-induced dynamics have been studied, such as stochastic resonance (25, 26), noise-induced order (27), consistency (8, 9), noise-induced synchronization in chaotic oscillators (28), excitable media (29), and periodic phase oscillators (30). For example, stochastic resonance (SR), which is a counterintuitive phenomenon where an optimal level of noise can enhance the responses of a nonlinear system to weak inputs, has been shown to play functional roles in animal and human brains. SR has been observed to improve sensitivity to sensory inputs in crayfish mechanoreceptor cells (31), paddle fish feeding behavior (32), and human visuo-motor coupling (33, 34). Apart from SR, there is only sparse evidence of other noise-induced phenomena associated with information processing in the brain, besides the above-mentioned single cell-level reliability studies (5, 6). We hereby add another piece of evidence that noise-induced dynamics in macroscopic brain responses are observable and robust, and that they can play roles in information processing in the brain. It should also be noted that the consistency nature observed in the current study provides evidence that macroscopic cortical dynamics exhibit a reservoir computing property in noise-induced dynamics.

Our study provides a novel manipulative method to probe the neural dynamics of the human brain. One of the efficient ways to probe the internal state or structure of such a complex system is to “ping” the system with a pulse input. Such perturbation approaches have been widely undertaken in physical, chemical, and biological systems, including the human brain. For example, transcranial magnetic stimulation (TMS) with concurrent EEG recording is used to probe human brain features such as effective connectivity (35) and area-dependent natural frequencies (36). We also reported a repetitive TMS paradigm to probe phase-amplitude coupling with concurrent EEG recording (37). In addition, Wolff et al. recently reported a new impulse response paradigm to probe the instantaneous hidden neural state associated with working memory (38). The frequency tagging method has also been of wide use for probing processing-related areas in the visual system (39). These studies indicate that perturbation approaches provide versatile tools for understanding the dynamical features of the brain. However, as far as we know, no prior studies have used noisy continuous inputs to probe the dynamical nature of the human brain. Thus, this study is novel in that it provides empirical evidence that noisy inputs can probe the dynamics of the brain, irrespective of differences in its initial condition, in a system- and input-dependent way.

We speculate that our dynamical method takes advantage of EEG-level neural dynamics, which show higher temporal precision than fMRI, despite the low spatial resolution. Recent human functional magnetic resonance imaging (fMRI) studies show that functional connectivity profiles during rest or task performance work well as a “fingerprint” to identify individuals (40). Although we used a high-density 63-channel EEG recording system, EEG is subject to the limitation of its poor spatial resolution in comparison with fMRI. It is therefore surprising that, despite the potential misalignment of electrode locations and different physiological conditions over sessions, the results were stable over periods of days. The current study provides more evidence that human EEG dynamics reveal individuality and inter-individual variations, building on our previous study on the visualization of resting-state EEG dynamics based on manifold learning (41).

The decoding of stable information from fluctuating brain responses against identical inputs is very important for achieving brain–machine interfaces (BMIs). Instability of brain signals is always an issue for practical BMIs because brain signals fluctuate even within individuals as they are disturbed by task-irrelevant brain activity (42). Therefore, consistency and the echo-state property are key factors in overcoming such instability in brain signals and facilitating BMIs. In fact, an EEG fingerprint study utilized event-related potentials (ERPs) as individual signatures (43). However, ERP-based methods need a number of trials (50–100 times) to allow a clear ERP to be attained, as task-irrelevant activity is cancelled out by the ensemble averaging of noisy brain signals. By contrast, in our consistency-based method, a 5.5 s single trial is enough to obtain excellent performance in individual verification if there is a target “database” to check. This advantage of our method over conventional ERP-based methods could facilitate versatile applications in the BMI field if we optimize the methods in terms of the experimental paradigms. For instance, the idea of physical reservoir computing (44) utilizing the consistency property of human EEG dynamics will facilitate future BMI applications.

## Materials and Methods

### Experimental Design

#### Participants

A total of 150 volunteers (74 males, 76 females; mean age, 24.2 years, standard deviation [SD] 5.2) with normal or corrected-to-normal vision participated in this study after giving informed consent. Among them, 130 participants underwent an experiment with noisy flickering stimuli and 20 participants underwent an experiment with periodic and noisy flickering stimuli. The study was approved by the ethics committee of RIKEN.

#### Apparatus and stimuli

The visual stimuli were presented on an LCD monitor (BenQ XL2420, BenQ corporation, Taipei, Taiwan; refresh rate: 60 Hz; resolution: 1024 × 768; black background luminance: 0.06 cd/m2) using NBS Presentation (Neurobehavioral Systems Inc., Berkeley, CA, USA). A chin rest was used to maintain the head position at a distance of 100 cm from the monitor.

The checkerboard stimuli (visual angle, 8.86 degrees) consisted of a 7 by 7 grid of gray squares with their gray levels being temporally modulated between black and white (luminance of black: 0.06 cd/m2, white: 48.8 cd/m2). The stimuli consisted of a fixation cross, a static checkerboard (2.5 s), and a noisy flickering checkerboard (5.5 s). The noisy stimuli were presented on a black background at 30 Hz, flickering at every two consecutive frames. The gray-level contrast between adjacent squares was modulated according to Gaussian white noise with a mean of 0 and SD set to 16, 32, 48, 64, or 80 gray levels. The mean gray level of the squares was set to 127. Two different temporal presentations of five different levels of Gaussian white noise were given to the participants. The correlation coefficient between the two different presentations of noisy visual stimuli was around zero. The contrast of static stimuli was set to 128 gray levels for all noise level conditions (n = 110), or to 16, 32, 48, 66, and 82 for the noise conditions with SDs of 16, 32, 48, 64, and 80, respectively (n = 20). Visual stimuli were generated using MATLAB (The MathWorks, Inc., Natick, MA, USA), and the visual experiment was implemented using Presentation software (Neurobehavioral Systems, Inc., Albany, CA, USA). Among the 130 participants, 32 were tested in a follow-up session after an interval of a few months (mean ± SD, 101.3 ± 60.2 days; range, 29–245 days).

Twenty additional participants were presented with periodic stimuli that flickered at four distinct frequencies (3.75, 5, 7.5, and 15 Hz) at a set intensity (lighter squares set to a gray level of 25, and darker squares to 229, contrast set to 204 gray levels), in addition to the highest level (SD 80) of noisy flickering.

#### EEG acquisition and preprocessing

EEG signals were measured using an EEG amplifier (BrainAmp MR+, Brain Products GmbH, Gilching, Germany) with a 63-channel EEG cap (Easycap, EASYCAP GmbH, Herrsching, Germany) at a sampling rate of 1000 Hz. The online lower and higher cutoff frequencies of the EEG amplifier were set to 0.016 Hz and 250 Hz, respectively.

Electrodes were positioned according to the international 10/10 system with a left earlobe reference and AFz as a ground electrode. EEG signals were offline re-referenced to the average of the left and right earlobe references and bandpass filtered between 3 and 80 Hz using a finite impulse response filter. In addition to the visual experiment, 3 minutes of EEG signals were measured while participants rested with their eyes closed. The resting EEG data were divided into 5.5 s epochs and used as noise SD 0 data. Single-trial EEG epochs were extracted around the visual signal presentation and from the 3-minute resting period, and an ICA-based automatic artifact removal procedure (ADJUST) was used to remove generic artifacts, eye movements, and blink-related artifacts (45). Next, the artifact-removed EEG signals *X*(*t*) were standardized between −1 and 1 using a hyperbolic tangent function (tanh)

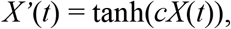

where *c* is set to 0.01.

All analyses were performed in MATLAB (The MathWorks Inc., USA) using custom-written scripts, ADJUST (45) and EEGLAB (46).

#### Intra and inter-individual CCA analysis

The EEG data recorded during the noise presentation consisted of 63 channels and 5500 samples per channel. There were 196 (= 14×14 trial pairs) combinations of pairwise EEG datasets between the different presentations of visual noise, and 182 (= 2 noise presentations ×_14_C_2_ trial pairs) combinations of datasets between identical presentations of visual noise. We applied CCA between these pairwise EEG datasets and evaluated canonical correlations across all possible trial-pair combinations. The canonical variates consisted of linear combinations of EEG signals using the 1st eigenvector of the CCA as weight coefficients, which indicated the maximal correlations between two EEG trials consisting of 63 channels and 5500 samples per channel. After obtaining canonical variates from pairwise EEG datasets, we computed the L1 norm between the two 1st canonical variates, which were 5500-point time-series data.

In the case of inter-individual analysis, there were 196 (= 14× 14 trial pairs) combinations of trial pairwise EEG datasets between a pair of individuals when viewing the same presentations of visual noise. In total, there were 8385 (= _130_ C_2_) participant pairs for 130 participants.

#### Visual inputs-EEG CCA analysis

The relationship between visual inputs and EEG responses was also analyzed by conducting CCA between the time-series data of the visual noise contrast (1 × 5500 time points) and the EEG signals (63 channels × 5500 time points). Specifically, we checked whether flipping the relationship between visual noise and EEG signals changed the canonical correlations. In this case, there were 14 × 2 combinations for correct relations of visual noise and EEG data within each individual, and 14 × 2 combinations for flipped relations. Notably, group analyses showed no significant change in canonical correlations (Wilcoxon signed-rank test, two-sided, n.s. p = 0.6864) after flipping the visual noise to EEG signal combinations. These results indicate that the EEG responses are not merely linearly transformed sequences of visual inputs.

#### t-SNE visualization

t-SNE analysis was conducted using a MATLAB t-SNE function. The two-dimensional Mahalanobis distances between 162 individuals (1st recording *n* = 130, 2nd recording *n* = 32) were Box-Cox transformed and used as an individual feature for the t-SNE analysis.

#### Statistical Analysis

The statistical significance level was set at p < 0.05. Statistical analyses were conducted using the MATLAB Statistics and Machine learning toolbox.

## Acknowledgments

We thank Mr. Motonobu Fujioka and Mr. Shunta Yoshimura for help with the data analysis. We thank Dr. Mitsuo Kawato for helpful discussion and comments.

## Author contributions

KK contributed to conceptualization, formal analysis, funding acquisition, investigation, methodology, project administration, resources, software, supervision, validation, visualization, writing – original draft, writing – review and editing. TS contributed to formal analysis, software. YM contributed to investigation. HS contributed to conceptualization, formal analysis, funding acquisition, investigation, methodology, software, validation, visualization, writing – review and editing.

## Data and materials availability

All data, code, and materials used in the analysis are available upon reasonable request.

## Supplementary Materials

### Power spectrum of the canonical variates

The participant-averaged power spectrum was computed for the 1st canonical variates for EEG trial pairs for participants viewing the same noise presentations (n = 130). Although there was a weak peak at 30 Hz corresponding with the frame rate of the noisy checkerboards, fast Fourier transform analyses revealed that the 1st canonical variate extracted from the paired EEG epochs for the same visual noise presentations showed several peaks in the 4–8 Hz theta band, with these being more prominent at high noise intensity conditions (Fig. S1).

**Figure S1.**
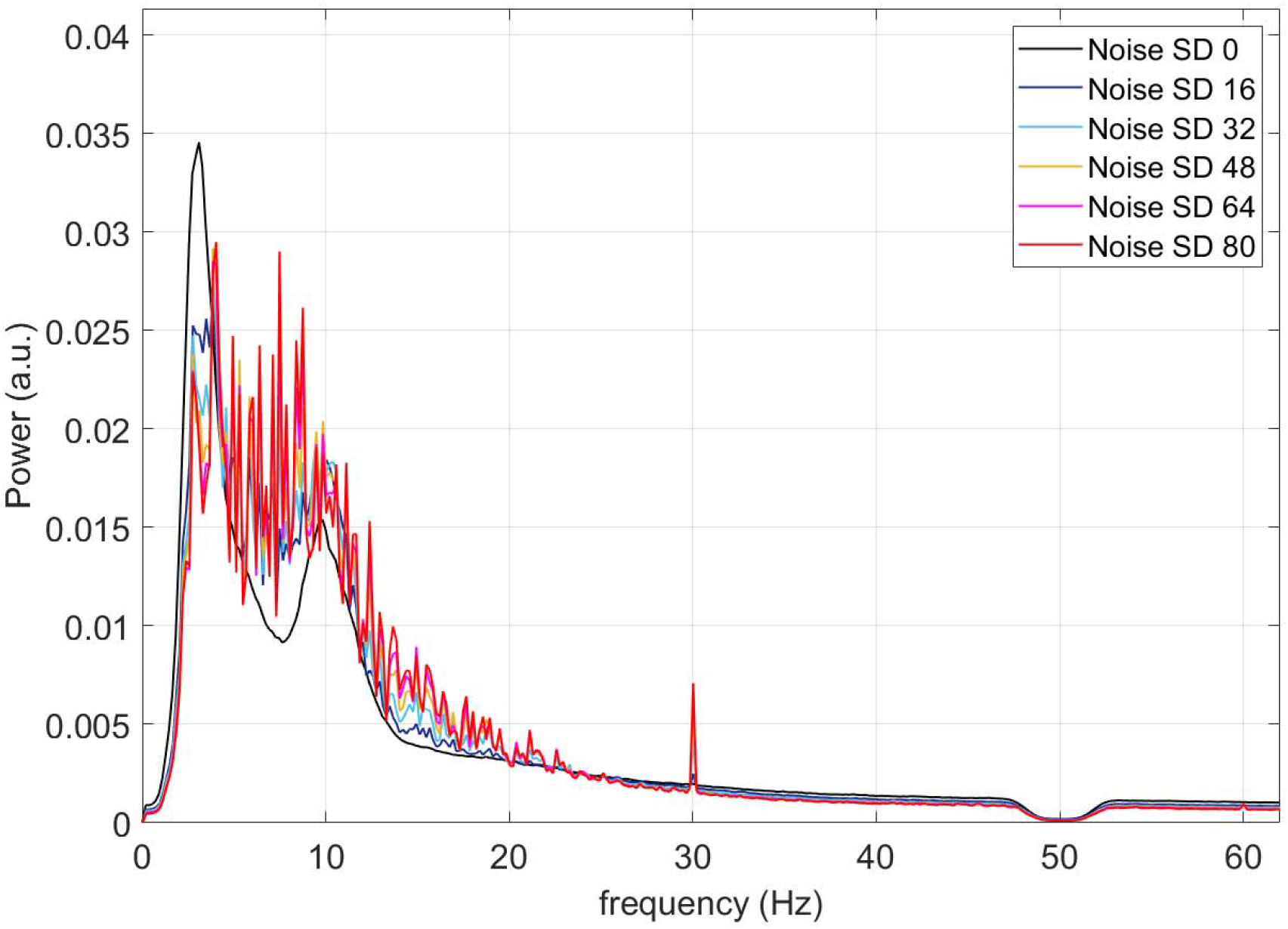
Power spectrum of the 1st canonical variates for each noise intensity.

### Results for periodic flickering

In 20 additional participants, the degree of inter-individual separation was compared between noisy and periodic inputs (3.75 Hz, 5.0 Hz, 7.5 Hz, 15.0 Hz) using the Mahalanobis distance and SVM with LOOCV. The EEG Mahalanobis distance between different individuals in the case of noisy flickering (SD 80) was significantly higher than that with periodic flickering (such as the SSVEP signals) and the resting condition (noise SD 0). There were significant effects of conditions on the Mahalanobis distance (Friedman test, Fr (5, 945) = 481.6, p < 0.001). Post-hoc multiple comparison tests showed that noise with an SD of 80 showed higher Mahalanobis distances between individuals than did the other conditions (Wilcoxon signed-rank test, two-sided, Bonferroni-corrected, p < 0.001; Fig. S2 (A)).

The conditions also had significant effects on the classification accuracy (Friedman test, Fr (5, 945) = 410.2, p < 0.05). Post-hoc multiple comparison tests showed that noise with an SD of 80 showed a higher classification accuracy than the other three conditions (7.5 Hz, 15.0 Hz, SD 0; Wilcoxon signed-rank test, two-sided, Bonferroni-corrected, p < 0.05; Fig. S2 (B)).

### Power spectral of canonical variates

**Figure S2.**
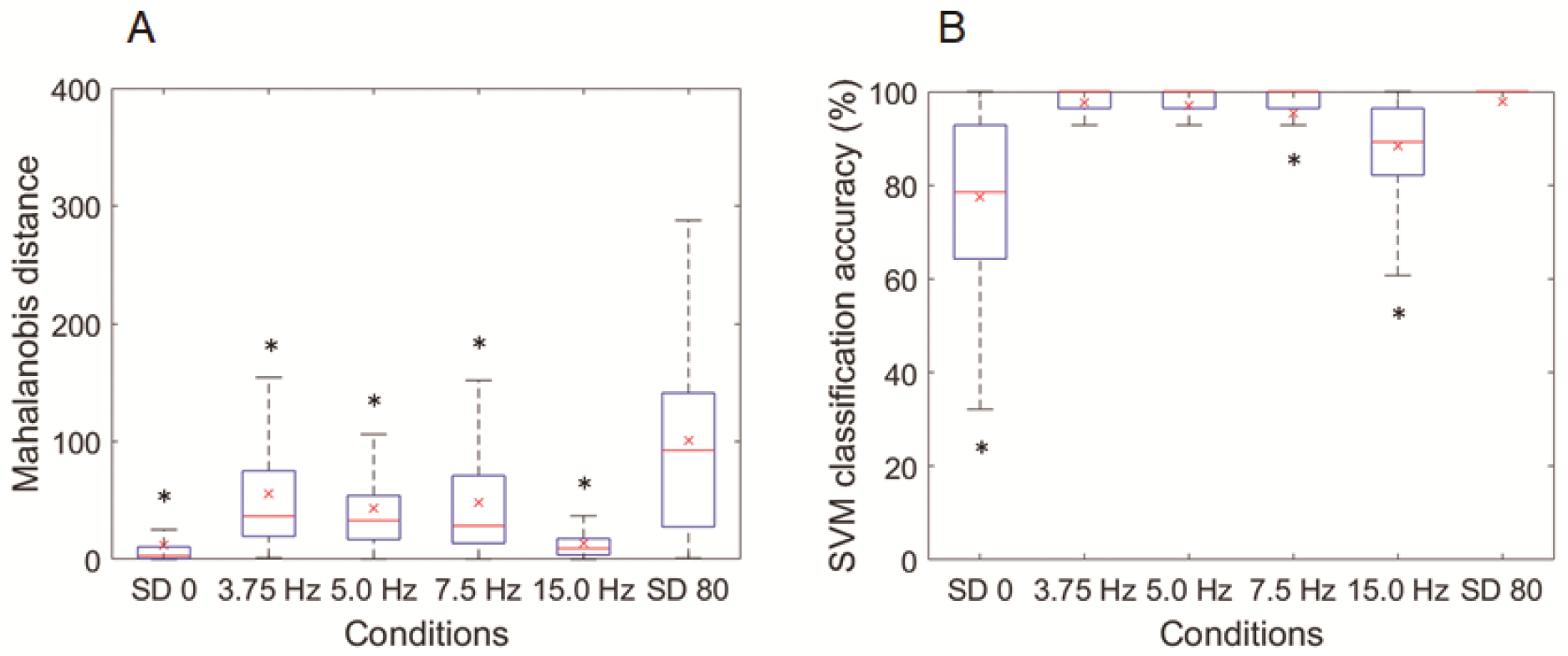
Comparison between noisy flickering and periodic flickering. (A) Group data for Mahalanobis distance between individuals (n = 190 [= _20_C_2_] participant pairs). (B) Group data for SVM LOOCV accuracy for classifying pairwise individuals (n = 190 [= _20_C_2_] participant pairs). Box plots in (A) and (B) show median, 25% and 75% quartiles (boxes), range (whiskers), and mean (x). Asterisks indicate significant differences between SD 80 and other conditions.

